# Covert attention to obstacles biases zebrafish escape direction

**DOI:** 10.1101/2022.04.14.488363

**Authors:** Hanna Zwaka, Olivia J McGinnis, Paula Pflitsch, Srishti Prabha, Vikash Mansinghka, Florian Engert, Andrew D Bolton

## Abstract

To study the evolutionary origins of object perception, we investigated whether a primitive vertebrate, the larval zebrafish, is sensitive to the presence of obstacles. The zebrafish, which has become a useful model to study brain-wide circuit dynamics, executes fast escape swims when in danger of predation. We posited that collisions with solid objects during escape would be maladaptive to the zebrafish, and therefore the direction of escape swims should be informed by the locations of barriers. To answer this question, we developed a novel closed-loop high-speed imaging rig outfitted with barriers of various qualities. Using this system, we show that when larval zebrafish escape in response to a non-directional vibrational stimulus, they use visual scene information to avoid collisions with obstacles. Our study demonstrates that fish compute absolute distance to obstacles, as distant barriers outside of collision range elicit less bias than nearby collidable barriers that occupy the same visual field. The computation of barrier features is covert, as the fish’s reaction to barriers during routine swimming does not predict that they will avoid barriers when escaping. Finally, through two-photon laser ablations, we suggest the presence of an excitatory input from the visual system to Mauthner cells in the brainstem escape network that is responsible for escape direction bias. We propose that zebrafish construct “object solidity” via an integrative visual computation that is more complex than retinal occupancy alone, suggesting a primitive understanding of object features and possibly the origins of a structured model of the physical world.

## Introduction

The ability of humans and animals to interact with the environment is mediated by our understanding of how the physical world works (i.e. “intuitive physics”; [4, 36]). There are two accounts on how information available in the world is transformed into action by the brain. In one account, evolution has established a mapping between patterns of neural activity on the sensors (e.g. the retina) and corresponding adaptive behaviors. Under this “reactive policy”, animals can be thought of as input-output machines resembling a Roomba [8], which flourishes within its limited environment and displays seemingly complex behaviors that are built up from simple rules. On the other hand, higher animals appear to demonstrate cognitive flexibility that suggests a structured model of the world: objects have three dimensions, identities, and properties that exist through time and space. Objects can be classified without visible action (e.g. “covert” attention), and our knowledge about the scene structure can be used to explicitly predict future states and outcomes (e.g. collisions). This type of predictive capacity likely arose as the environment become so complex that a fixed reactive policy for each situation would require almost infinite computational capacity. It remains an open question at what point in evolution this transition from reactive behavior to structured knowledge began to occur? Earlier studies had suggested that simple animals like chicks and frogs possess some aspects of physical cognition [10, 17, 34]; we therefore wondered whether an even simpler, neurally accessible organism, the larval zebrafish, would display behaviors that rely on the representation of objects as solid structures.

To this end, we examined the sensitivity of larval zebrafish to the locations of barriers during predator escape. In zebrafish, escapes from predators or artificial predator-mimicking stimuli are controlled by a well characterized population of neurons called the “brainstem escape network” (BEN) [13, 14, 29]. The BEN, containing the Mauthner neuron pair which is capable of evoking high-angle tail bends in response to startle, is experimentally accessible to genetic labeling and two-photon laser ablation. Critically, this circuit mediates a two alternative behavioral choice [19]: fish direct their escape trajectories to either the left or the right. However, the extremely short latency and ballistic nature of these movements make accidental collisions with surrounding objects a hazard. Therefore, it would be adaptive for the fish to generate a bias to their escape behaviors that directs them away from detected barriers.

Here we show that startled zebrafish reliably avoid collisions by directing their escapes away from barriers (i.e. if the barrier is on the left, they escape right and vice versa). We confirm that visual signals are necessary for the induction of the bias, since fish execute unbiased escapes and collide randomly with barriers in the dark. Keeping the retinal occupancy steady but varying physical size and distance of barriers, we demonstrate that fish are sensitive to the absolute distance of barriers and use a more complex mechanism to measure distance than visual occupancy alone. To account for this result, we propose plausible metrics that fish could use to compute distance out of a set of previously described methods used by animals and humans [28, 32]. Lastly, we suggest the presence of a circuit motif whereby visual information excites the Mauthner neuron responsible for escapes away from barriers. In summary, we propose that larval zebrafish implicitly represent barriers as solid objects with a precise 3D location and identity.

## Results

### Barrier Avoidance During Escape

We characterized the escape trajectories of 99 wild-type larval zebrafish (WIK) when presented with a non-directional startle stimulus. Fish were imaged at high speed (500 Hz) while swimming in a 12 cm diameter tank (Figure 1A). Custom computer vision software was designed to detect when fish entered a 12 mm wide circular zone located 4 cm from the edge of the tank. The location of the zone was unmarked and unknown to the fish. Upon zone entry, the detection software triggers a “tap” via an electromagnet that strikes the tank with a metal rod for 200 ms. This stimulus reliably evoked fast escapes at extremely short latency ([22], Figure 1A, right panel). The direction of escape in the absence of barriers was random (Figure 1B,C). We introduce two indices that describe the direction of escapes in our assay. First, Preference Index (PI) is (#Rightward Escapes - #Leftward Escapes) / (# Total Escapes). Calculating this metric for each fish in the absence of barriers yielded a median preference index of 0.0 and mean of -.03, indicating no average preference for one direction or the other (p = .503, 1 sample ttest population mean different from 0.0, Figure 1C). Individual fish are most commonly unbiased in their escape which is indicated by the highest density of the normal PI distribution centered around the 0.0 bin (Figure 1C). Figure 1B (left panel) shows the time course of escape trajectories following stimulus delivery, and the 2D histogram below illustrates the unbiased distribution of locations visited by the fish immediately after the tap.

**Figure 1.**
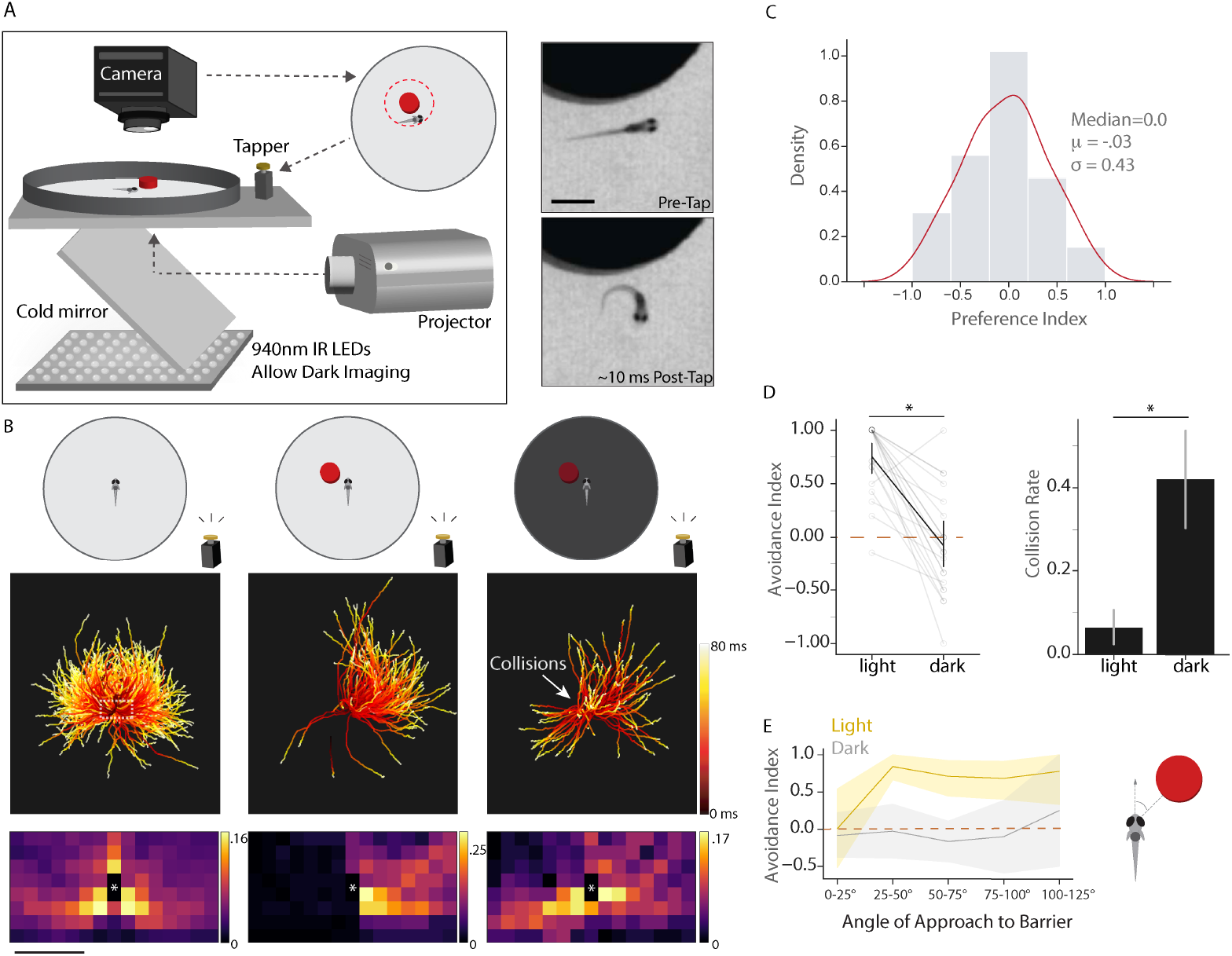
Zebrafish switch from randomly directed to biased escapes in the vicinity of barriers. A. Left: Overview of experimental setup. Right: Typical large-angle short-latency escape turn after tap delivery. B. Escape trajectories after tap induction, color coded to reflect time, are shown for 3 conditions (no barriers in visible light, barrier in visible light, barrier in the dark). Trajectories begin 20 ms after tap command and trajectories with barrier approaches to the right are reflected across the Y axis. 2D histograms depict probability of the escape trajectory passing through a spatial bin for each condition. White asterisk indicates starting position of the fish. Barriers are 12mm wide and 6mm high. Fish are tapped when they pass within 2 mm of the barrier. (N=99, n=967; N=22, n=128; N=21, n=102) C. Distribution of Preference Index for Left vs. Right escapes in no barrier conditions (N=99). D. Pairwise comparison of barrier avoidance index in fish tested in both the light and dark. Collision rate in light vs. dark. (same fish as B) E. Avoidance index per fish per window of approach angle to the barrier in the light (yellow) and in the dark (light grey). In the light, escapes are not biased in the frontal 25° of visual space but are significantly biased (95% CI above 0) for all other windows. Solid lines represent mean and shaded areas 95% CI. Scalebar in trajectory plots = 2.5 mm; Scalebar in 2D histograms = 1.25 mm. Area in dotted box in left B panel reflects area plotted in 2D histograms.

Each fish shown in the left panel of Figure 1B was also tested in an additional assay assessing the same behavior in the presence of barriers. To measure barrier avoidance, we introduce a second statistic, Barrier Avoidance Index (BAI), which is (# Escapes Away From Barrier - # Escapes Towards Barrier) / (# Total Escapes). For our initial barrier condition, we placed a 12 mm wide, 6 mm high red acrylic barriers into the tank at the same relative tap zone described above, triggering taps when fish passed within 2 mm of the barrier. The introduction of a barrier completely changed the distribution of escape directions. Fish significantly directed their escape trajectories away from the barrier (Figure 1B, center panel) with an average BAI of .78 (p = 2.9e-10). This bias away from barriers, however, disappeared if the same fish were tested in the dark (Figure 1B, right panel; p = .52). Pairwise comparisons of BAI for each fish in light and dark conditions are shown in Figure 1D, which also shows that the barrier collision rate significantly increases from 6% when fish can see the barrier to 42% in the dark (p = 7.32e-6, paired ttest). We therefore conclude that zebrafish use vision to detect barriers and reliably avoid collisions by converting their typically random escape directions into laterally biased trajectories.

One phenomenon observed during barrier trials that points to the fish’s behavioral algorithm shows that BAI depends on the angle of barrier approach. In the dark, escapes were unbiased regardless of the angle of approach (Figure 1E). However, in light conditions, if fish approach barriers head-on (0°-25° on either side), their escape trajectories are unbiased and resemble trajectories in dark conditions (bootstrap 95% confidence interval contains 0.0 BAI, yellow line Figure 1E). This suggests that lateral movement of the barrier during approach may be involved in generating bias. In this study, we therefore exclude head-on barrier approaches (−25° to 25° angle of approach) when making comparisons between barrier conditions.

In order to investigate which features of barriers trigger avoidance, we performed the tap assay while varying the height, width, distance, and color of the barrier. BAI under all tested conditions is plotted in descending order of effect in Figure 2. Doubling the distance (4 mm) to the same sized barrier as in Figure 1 continued to bias escapes away from barriers (p = .007 population mean different from 0.0), but reduced the average BAI magnitude from .78 to .27 (p = 5.4e-5, two-sample independent ttest, Figure 2A). This reduction in BAI scaled directly with the predicted probability of colliding with the barrier (45% collision rate to 19%, Figure 2A top and Figure 2 legend), consistent with the hypothesis that zebrafish map barrier locations to avoid collisions. We next wondered whether the decreased *apparent* width and height of the barrier due to increased distance was responsible for this reduction. Doubling the height of the 4 mm distant barrier (12 mm high) did not significantly change BAI (.27 to .36, p = .48). However, doubling the width (24 mm wide) nearly restored BAI to original values (.27 to .67, p = .001, original value.78), initially suggesting that the amount of horizontal retinal occupancy may be a key factor in determining bias. We tested this idea by placing a barrier that occupied the exact same amount of the horizontal and vertical visual field as the 24 mm wide barrier but instead at 8 mm away, reducing the probability of collision to 0 (Figure 2 top panel). Interestingly, vertical and horizontal visual field occupancy was not the most important factor in biasing escapes. Although apparent size is identical between the two conditions (24 mm w, 6 mm h, 4 mm d vs. 48 mm w, 12 mm h, 8 mm d), distance was the determining factor in biasing escapes, as BAI dropped from .67 at 4 mm distance to .22 at 8 mm distance (p = .0002, two-sample independent ttest). Coupling this result to predicted collision statistics in Figure 2 suggests that the zebrafish’s escape strategy accounts for not only how far away barriers are in their environment, but how far their escape trajectories propel them in space. The .22 average BAI in distant barrier conditions shows that escapes do remain slightly biased from 0.0 (p = .037) if the barrier is 12 mm high, but if height is dropped to 6 mm, we encounter the only barrier condition where fish do not bias from (rightmost condition, Figure 2; BAI = .07, p = .68). Therefore, there appears to be a threshold vertical occupancy; a 6 mm high, 8 mm distant barrier occupies 35° of visual angle, suggesting the threshold for vertical occupancy is between 35° and 70°.

**Figure 2.**
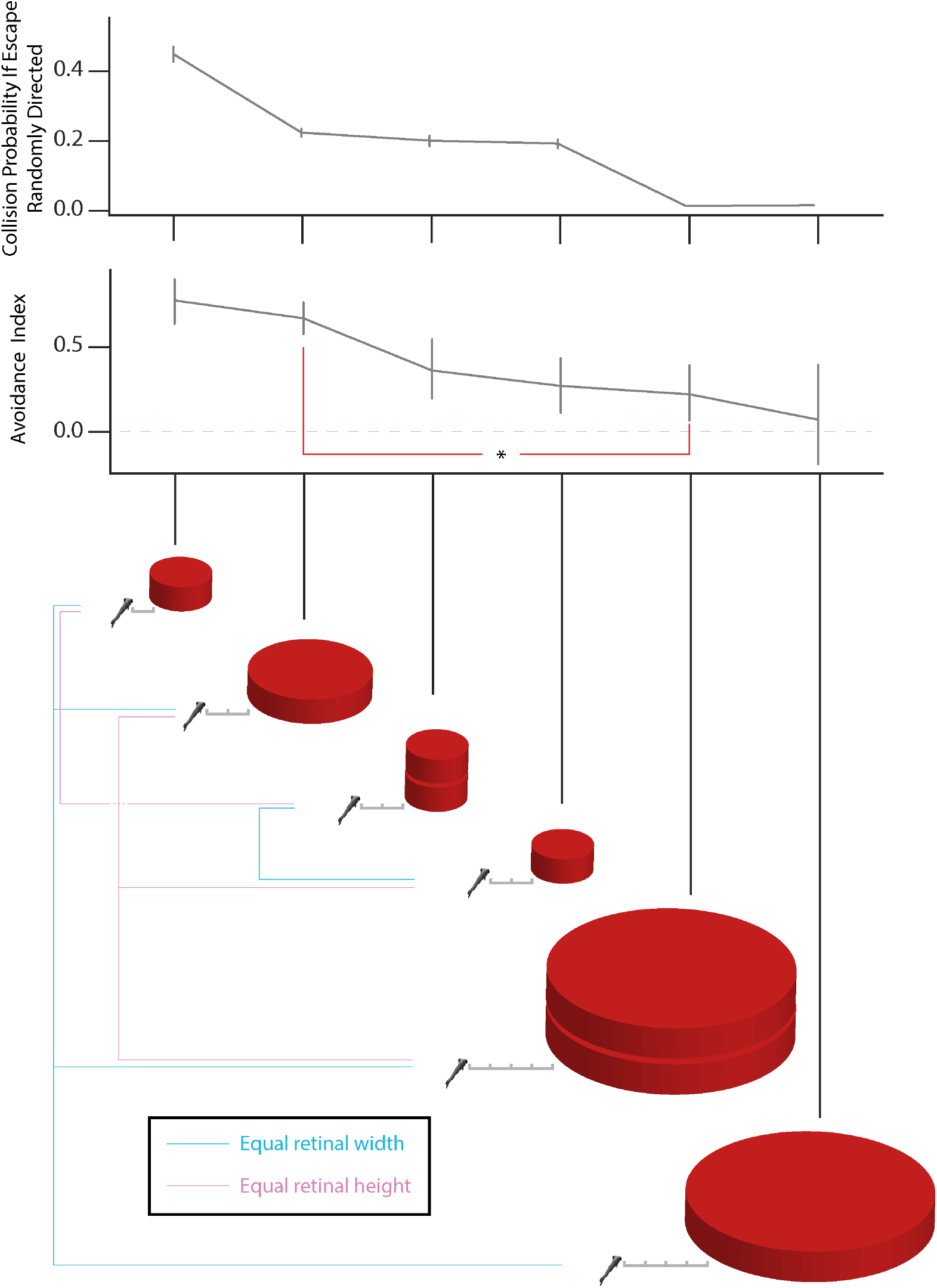
Barrier avoidance depends on barrier size and distance. Collision probability is calculated as follows: for each trial within a barrier condition, the heading angle of the fish and the angle to the barrier immediately preceding escape are stored. Each of the 967 trajectories in Figure 1B (no barrier), which reflect the escape behavior of unbiased fish, are then simulated from the stored starting conditions. Intersections between simulated control trajectories and barriers are recorded and divided by the total # of control trajectories to obtain a collision probability per trial. Collision probability therefore reflects how often an unbiased fish would collide with the barriers that subjects encountered during the different barrier conditions. BAI decreases in correlation with collision probability. BAI is also significantly different for barriers with the same retinal occupancy but different distance (red line). (N=22, 17, 16, 13, 12, 7; n=128, 150, 148, 122, 104, 56). Scalebar ticks in diagram = 2 mm

### Barriers Influence Routine Swimming

We next asked whether barrier avoidance during escapes was covertly pre-computed or whether fish overtly show evidence of barrier aversion before the tap stimulus arrives. 12 fish were allowed to freely swim in the tank with four 12 mm wide barriers placed according to Figure 3A. We find that fish are sensitive to barrier locations even in the absence of tap evoked escapes. In Figure 3A (left), the complete trajectory of a free swimming fish is displayed. Figure 3B (left) displays a density plot with the number of visits to each bin of space across all fish. The vicinity surrounding red barriers is the least visited region of the tank, suggesting that fish treat red barriers as aversive. We conclude that the fish performs covert avoidance of barriers regardless of its overt routine behavior.

**Figure 3.**
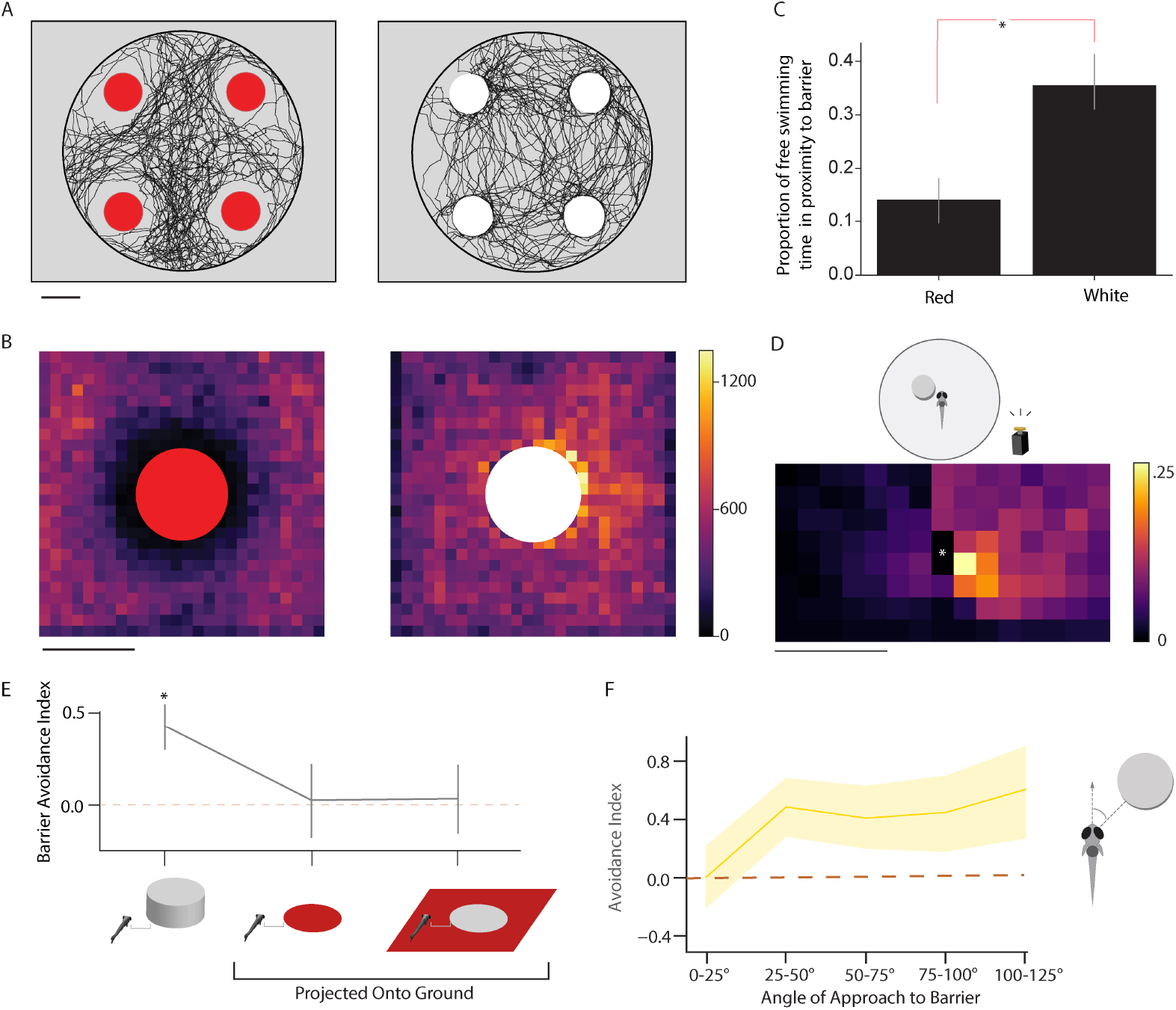
Red barriers are aversive and white attractive, but both are avoided during escape A. Example trajectories of a zebrafish freely swimming in the arena with four red barriers (left panel) or four white barriers (right panel; N= 6, 6). B. Total visits to each spatial bin surrounding red and white barriers. C. Proportion of swimming time in the vicinity (20 × 20 mm) of barriers is significantly different for red and white barriers. D. 2D histogram depicting probability of the escape trajectory passing through a spatial bin for white barrier conditions. White asterisk indicates starting position of the fish. (N=19, n=254) E. Average barrier avoidance index for white physical barriers and barriers projected onto the floor (N=19, 20, 20; n=254, 160, 146) F. Avoidance index for different angles of approach to the white barrier in the light (yellow). As with red barriers, escapes are not biased in the frontal 25° of visual space. Solid lines represent mean and shaded areas 95% CI via bootstrap. Scalebar in A, B = 12 mm; Scalebar in D = 1.25 mm

Although inherent aversion could be a strategy for avoiding collisions during escape, we wondered if we could find a barrier condition that could dissociate avoidance during escapes from aversion during free swimming. We found that replacing red barriers with white barriers of the exact same size completely inverted preference during free swimming (Figure 3A-E, p = .0002, ttest for % time spent in the 20 × 20 mm region surrounding red vs. white barriers). Instead of avoiding barriers, the zebrafish’s trajectories reflect an attraction to white barrier locations, and spatial bins bordering the white barrier were the most frequently visited (Figure 3A+B, right panel). Remarkably, zebrafish continued to treat white barriers as collidable objects during escapes, significantly biasing their escape turns away from barriers (Figure 3E, BAI = .43, p = 3.05e-6).

Like red barriers, escapes away from white barriers are not biased in the frontal 25° of visual space, and spatial bins on the side opposite the barrier are the most commonly visited during escapes (Figure 3F), suggesting the same bias mechanism. Although fish avoid white barriers two out of every three escapes, this nonetheless reflects a significant decrease in BAI from red conditions (.78 to .43, p = .0008), where barriers are darker than the floor of the tank. We therefore asked whether phototaxis, wherein larval fish swim towards the brighter visual hemifield, could play a minor role in biasing escape direction. Zebrafish, however, did not bias their escapes if tapped within 2 mm of a red dot projected onto the bottom of the tank nor a white dot projected onto a red background (Figure 3 E right side, BAI = .028, .037; p = .79, .71; dots are equal width of barriers). This suggests that color comparison between the left and right hemifield does not itself play a role in escape, and that barriers must have height to be considered an obstacle. We surmise that the small difference in BAI between red and white conditions is due to a modest interaction between covert avoidance and overt attraction.

### Mauthner Cell Ablations Reveal a Biasing Circuit Motif

We next performed an initial circuit analysis to determine how the visual system influences the Brainstem Escape Network (BEN), which mediates two-alternative behavioral choice during escapes and contains the commonly-known Mauthner neurons [19]. Mauthner cells, along with their homologs MiD2cm and MiD3cm, play a prominent role in evoking short latency, high amplitude escape turns in fish [14].

Spiking in the right Mauthner neuron typically results in escapes to the left, while left Mauthner spiking results in escapes to the right. However, homologs can also control escape features even in the absence of Mauthner cell activity [22, 29]. We wondered how visual information about barrier location impinges upon this complex circuit. Critically, Mauthner cells receive modulatory inputs from throughout the brain that are conspicuously positioned for biasing escape direction. Since 97% of the taps evoked in our barrier dataset (Figure 2) resulted in rapid, large amplitude escape turns typically attributed to Mauthner cells (mean 104.4° cumulative tail angle at 11.06 ms post-tap, Supplementary Figure 1), we surmised that barrier avoidance is accomplished via lateralized Mauthner modulation.

There are two ways for visual information to influence the Mauthners (Figure 4A): visual activity could either inhibit the Mauthner neuron that pulls the fish toward the barrier or excite the neuron that takes the fish away from the barrier. To uncover whether either of these motifs exist in the fish brain, we fluorescently labelled Mauthner neurons using a custom Gal4 line and ablated a single Mauthner neuron per fish with a two-photon laser (see Methods, Figure 4B). As noted, escape direction is typically contralateral to the spiking Mauthner neuron, so we expected fish with ablations to preferentially escape away from the side of the intact neuron. In Figure 4C, Preference Index is calculated as (# Turns away from intact Mauthner - # Turns towards intact Mauthner / # Total Escape Turns). As expected, fish significantly biased their escapes away from the intact Mauthner side when tapped in the absence of barriers (mean PI away from intact Mauthner = .405, p = .00018; see barrier-less conditions in Figure 4C). This level of bias is in alignment with previous Mauthner ablation experiments [22]. When barriers were added to the tank, taps executed when fish approached barriers contralateral to the intact Mauthner cell continued to induce escape *away from* the intact neuron (and resultingly, *towards* barriers) at the same frequency as barrier-less conditions, showing a complete loss in the ability to bias escapes away from barriers (mean PI away from intact Mauthner = .372, p = .77, two-sample paired ttest vs no barrier, Figure 4 C, D). On the other hand, if fish approached barriers ipsilateral to the intact Mauthner cell, their preference for escaping away from the intact neuron was actually facilitated (mean PI away from intact Mauthner = .622, p = .0237 two-sample paired ttest vs no barrier, Figure 4 C,D). Coupled together, these results favor the excitatory circuit architecture proposed in Figure 4A. The intact Mauthner neuron does not appear to be inhibited by the perception of a contralateral barrier, shown by the continued escape towards barriers. The intact Mauthner neuron instead undergoes an excitatory bias during perception of a barrier on the ipsilateral side, enhancing bias up to the levels observed in Figure 2 in the strongest biasing conditions (Figure 4E).

**Figure 4.**
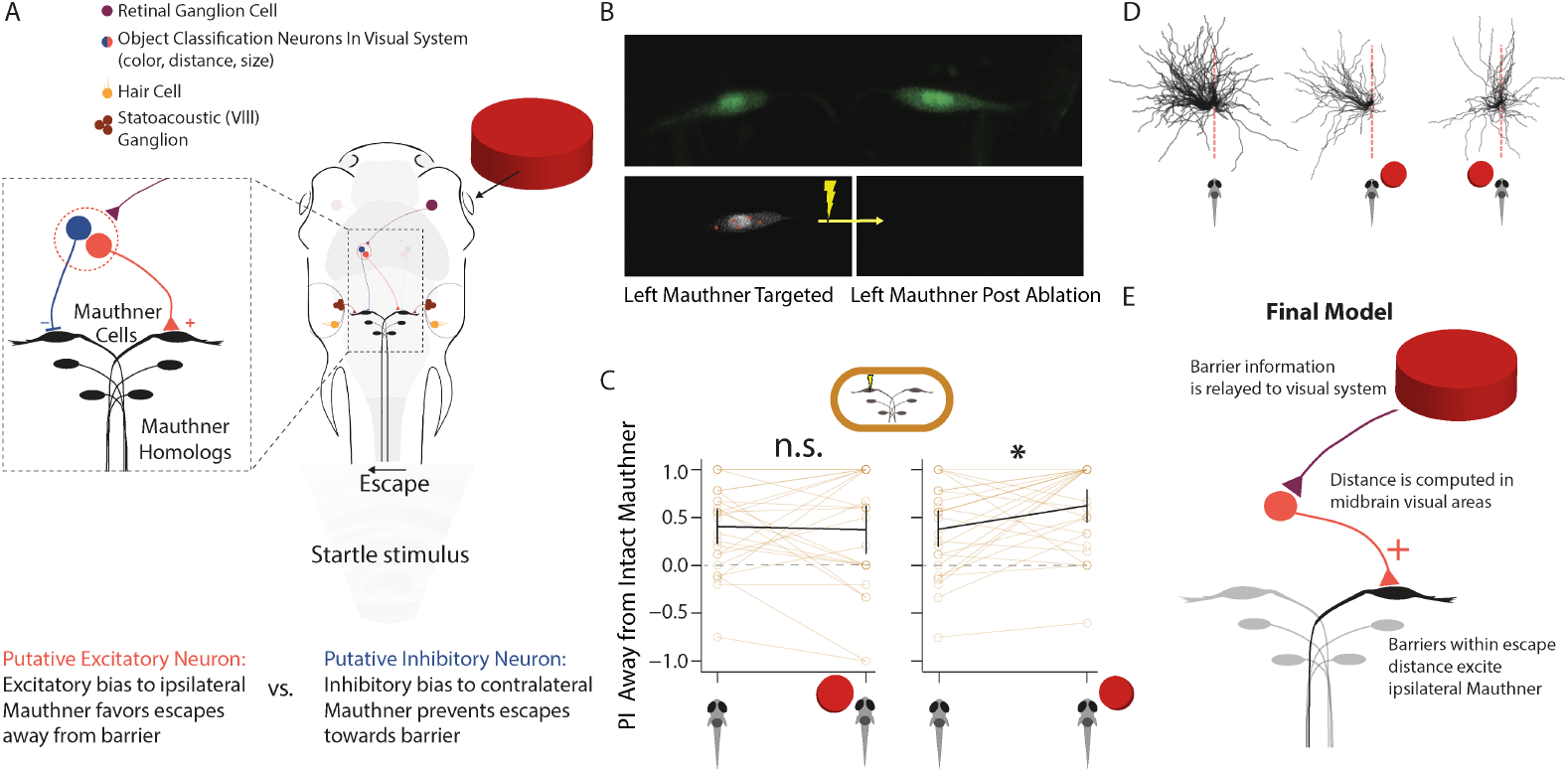
Excitatory visual input into ipsilateral Mauthner cell biases escapes. A. Two putative models can bias escapes away from a barrier: excitatory input into the ipsilateral Mauthner cell would enhance escapes away from a barrier or inhibitory input into the contralateral Mauthner cell would prevent escapes towards a barrier. B. Two-photon image of both Mauthner cells (upper panel) in our custom Gal4 line. Left Mauthner cell pre ablation showing our UV laser targeting protocol for Mauthner destruction. Same cell post ablation. C. PI for escapes away from intact Mauthner in control and barrier conditions. (N=26, n=275 control, 192 barrier) D. Individual escape directions for animals with the left Mauthner cell ablated. No barrier present with ablated right Mauthner cell leads to an escape bias to the left. Escapes with barrier on the right shows a strong bias to the left away from the barrier. A barrier on the left leads to a biased escape to the left in direction of the barrier, comparable to an escape with no barrier present with right ablated Mauthner cell. E. Final model after integration of Mauthner cell ablation experiments reveals excitatory input into ipsilateral Mauthner cell which favors escapes away from barriers.

## Discussion

Constructing intelligent behavior by combining simpler sensorimotor modules has become a recurring theme in neuroethology [2, 11] and artificial intelligence research [7, 8, 27]. Human-like intuitive physics is likely built upon the visuomotor neural circuits that have allowed animals to flourish in a complex physical world for eons without the benefit of explicit physical reasoning [8, 34]. In this study, we investigated the foundations of solid object perception by asking whether larval zebrafish are sensitive to the presence of obstacles during fast escape swims. We discovered that when zebrafish are startled in the vicinity of barriers, they alter their normal pattern of unbiased escape direction and instead consistently bias their escape swims away from collidable objects (Figure 1). This decision depends on barrier height and width, but most strongly on computed distance to barriers. Moreover, the magnitude of escape bias correlates well with the probability of collision (Figure 2). Using two-photon laser ablations, we imply the existence of an excitatory input to the Mauthner Neuron from the visual system that mediates escape bias. We suggest that the larval zebrafish embodies the concept of “object solidity” via connections from barrier detection modules to evolutionarily ancient predator escape circuits [25].

Biasing escape direction near obstacles is the first case of covert scene understanding that has been revealed in the larval zebrafish. By “covert”, we mean that the zebrafish has performed computations of barrier identification, size, and distance without explicitly showing it in its behavior. Only when evoking startle responses can we observe that the fish was attending the barrier. One could argue that our routine swimming experiments (Figure 3) show that the animals are overtly attending the differently colored barriers: red barriers are avoided while white barriers are approached. However, both barriers are avoided during escapes, suggesting that behavior in the moments preceding the escape does not predict barrier avoidance. Relatedly, the location of an adult goldfish within its tank in the moments preceding a predator stimulus modifies the fish’s ballistic escape direction; detection of tank walls as impermeable obstacles before escape initiation would be consistent with our findings [13]. Frogs similarly inform their choice of jumping direction according to the location of static objects [17]. Critically, the non-cortical brain regions that fish and frogs possess have been shown to play a role in covert computations of object identity in an array of animals [21, 23]. In frogs, “attention units” in the optic tectum respond to subtle movements of prey-like stimuli that do not immediately prompt hunting behavior. These subthreshold stimuli instead facilitate future prey striking behavior if the same prey object is presented seconds later. The neural correlate of this facilitation manifests when attentional units continue to respond seconds after the original stimulus has stopped moving. This study reflects the current idea that the midbrain covertly maps the location and identity of stimuli of interest in the visual scene [16], [20]. We classify this mapping as “implicit” scene understanding: object identification (e.g. “prey”, “barrier”) and location have been covertly computed for future behavioral goals.

Robust 3D vision requires accurate depth estimates for all items in a scene. Our results show that zebrafish larvae compute absolute distance to barriers, since bias to nearby barriers is significantly stronger than to distant barriers that occupy the same retinal area (Figure 2). Distance computation, in fact, has now emerged as an essential feature of both the zebrafish’s escape and prey capture behaviors [6], [30]. Previous studies have emphasized the role of stimulus “size”, measured by total retinal occupancy, in decoding stimulus identity [3]. However, total retinal occupancy, as shown here, fails to unambiguously convey the depth of a stimulus, and accordingly, its behavioral relevance. Depth must be *computed* from the 2D retinal representation – it does not come for “free” with any known visual map (e.g. an equal-sized neural representation on the optic tectum map could be a close prey or a distant barrier). This point is emphasized by previous studies examining the response of zebrafish to live prey and to different-sized moving dots. In prey capture, when a fish strikes at a 300 micron-wide prey item, the prey is so close (avg. 870 + 180 microns) that it occupies 16°-23° of visual space [6]. Moving dots occupying this amount of visual space induce the strongest possible *avoidance* in previous studies [3, 5]. Therefore, our study and previous results suggest that integration of multiple features, including contextual evidence, must be at play (see below) and the zebrafish’s capacity for scene understanding is greater than previously appreciated.

Interestingly, AI approaches seeking to replicate the brain’s ability to interpret 3D scenes almost exclusively rely on depth sensing cameras. This avoids the complexity that comes with inferring object locations from a 2D layer of photoreceptors that is itself incapable of pinpointing the depth of a light source. Studying biological distance sensing in simple animals could provide insights into how scene interpretation is so effortless for humans [18] yet difficult for computers. In fact, elegant studies in insects and amphibians have already painted a complex picture of how animals compute object distance, suggesting that dynamic lens accommodation and motion cues play important roles [24, 32, 35]. In addition to motion parallax observed in bees and flies, some insects like locusts and the praying mantis purposefully move their heads from side to side in order to gain distance information from self-motion [31, 33, 37]. We suggest that distance computations used by humans could be informed by these lower species because clinically blind humans with lesions in higher order visual areas can still navigate around obstacles without conscious awareness [15]. Similar results were obtained for reaching tasks around barriers in patients with temporoparietal damage, indicating that higher order visual awareness is not necessary for barrier avoidance during reaching [26].

The precise mechanism that larval zebrafish use to asses object distance remains unknown; however, brains use up to 13 methods to estimate distance (Figure 5) [28]. During a swim bout towards a barrier, the barrier will appear to expand, move laterally across the visual scene, and change focal plane, all in proportion to barrier distance when the swim began. Each distance calculation method comes with failure modes, but failures of different methods rarely co-occur. This allows thousands of accurate distance computations per day without crashing into walls, tripping over objects, or mistaking distant humans for nearby ants. Because barrier approaches that are “head on” fail to evoke a bias in our experiments (Figure 1E), we surmise that the amount of lateral movement across the scene (i.e. “motion parallax”) is a key component of distance computation, and that this variable is integrated with total amount of occupied vertical and lateral retina. Importantly, the window of visual space occupied by the barrier during “head-on” approach encapsulates the binocular zone [5], making it unlikely that fish use stereopsis for barrier avoidance. Future research should address what happens when distance metrics are forced to artificially conflict during virtual reality – how does the brain balance information from several channels that could each disagree?

**Figure 5.**
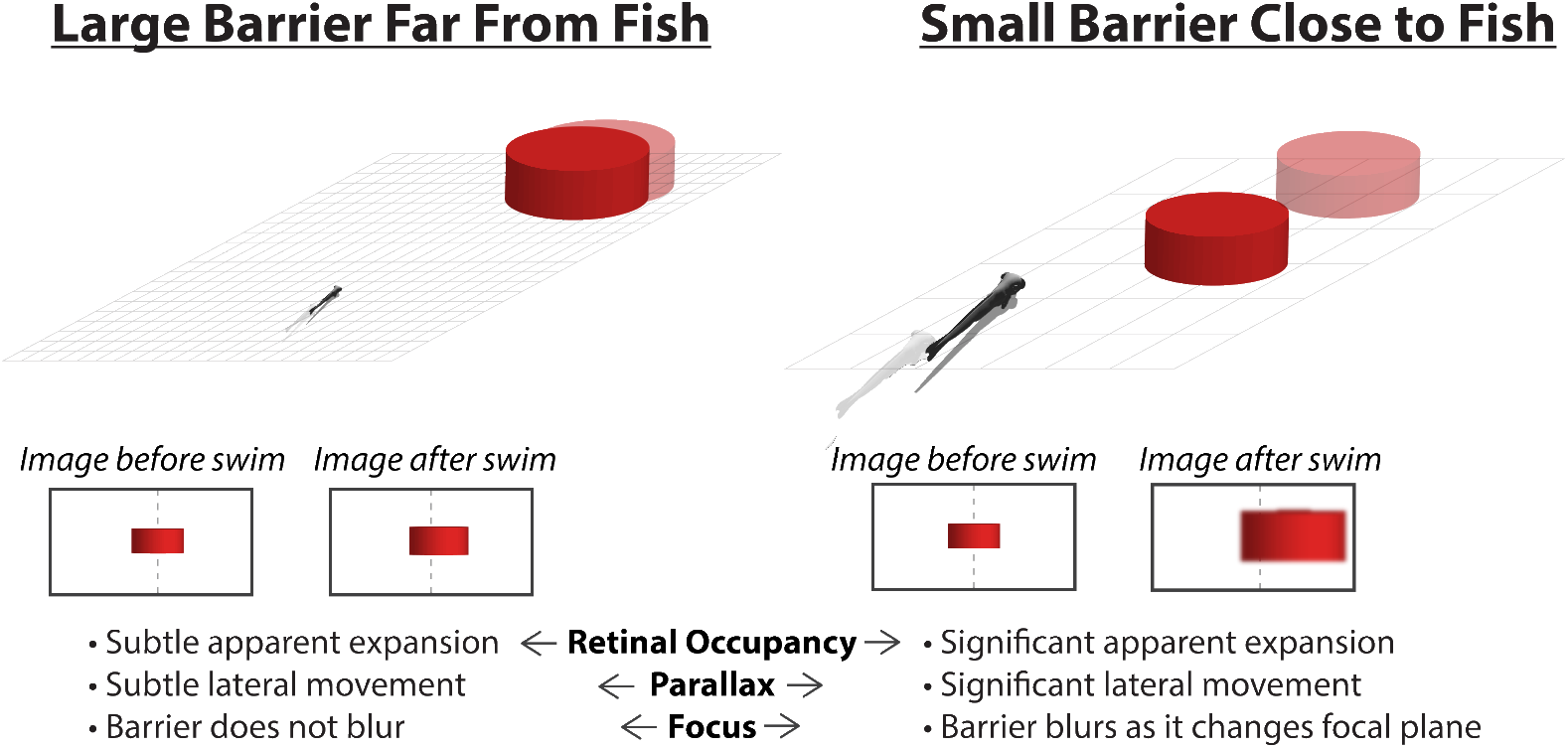
Three candidate mechanisms for distance computation in our assay.

As in our previous work [6], we propose that the fish’s ability to perceive distance is intertwined with its own body constraints. Zebrafish avoid barriers at distances that are within its escape swim limits (Figure 2), which resembles the idea of embodied cognition [25]; in other words, the fish’s escape system “knows” how far it displaces the body. Further evidence for embodied knowledge comes from the fish’s differential response to doubling the height vs. width of nearby barriers (Figure 2). Escape bias is significantly more sensitive to width changes, suggesting that the fish’s escape trajectory is likely to be more variable in the horizontal versus vertical plane. Future studies should address whether this is in fact the case, which would further support the idea that the co-evolution of sensory and motor systems should be considered during studies of perception.

We suggest that some features of the fish’s obstacle avoidance behavior (i.e. computation of absolute distance, covert attention) may suggest the presence of a structured model for physical cognition in the brain, or a primitive “physics engine” [4, 36]. However, our results do not exclude the possibility that the zebrafish is simply approximating a physics engine using bottom up feature detection combined with the perception of its own body thrust and the resultant motion of obstacles. Yet, the behavioral strategy executed by the fish in our study is more complex than a simple reactive strategy that translates retinal pixels and current behavioral state into resultant behaviors (e.g. optomotor response [1], phototaxis [9]); nevertheless, it may still fall short of the bar required for assigning “cognition” to the fish. How would future experiments disambiguate a complex bottom-up strategy from a structured cognitive model of objects in the world? One strategy would be to present fish embedded in a virtual reality setting with controlled visual stimuli that would disambiguate objects to a human or Bayesian observer. For example, if presented with a stimulus detected as a large distant barrier due to parallax, will the fish still approach a close by prey stimulus tuned to the specs (retinal size, angular speed) of a paramecium if the prey passes behind the barrier? Passing behind the barrier would suggest that the virtual prey is not a nearby prey at all, but a faster moving, larger object that is further away than the large barrier. If the fish pursues this stimulus as prey, a more reaction-based, sensorimotor intelligence in the spirit of Brooks would be assumed [8].

Lastly, with respect to the neural mechanism we hypothesize, ablating one Mauthner did not lead to an abolishment of escape towards the ipislateral side of the ablation. This was expected from previous studies on Mauthner ablation [22], and likely emphasizes the capability of Mauthner homologs to compensate for Mauthner loss. Importantly, touch stimulation of the head to induce escapes is the most effective means of recruiting Mauthner homologs, and leads to a more vigorous escape swim [29]. Our non-directional tap stimulus likely induces simultaneous head and tail stimulation, recruiting Mauthner homologs and allowing a bilateral competition between all BEN components across hemispheres. Our final hypothesis entails an excitatory input to the ipsilateral Mauthner cell that biases escapes away from barriers (Figure 4). In future experiments we plan on functionally dissecting this excitatory input which we hypothesize to come from the tectum via spiral fiber neurons [22]. Importantly, the type of elegant circuit analysis performed in previous studies of the zebrafish BEN [19, 22] is not yet possible for our behavior of interest using current technologies. We hypothesize that the fish requires motion parallax to estimate barrier distance, which necessitates a comparison between the motion of the barrier and the fish’s own motion vector (Figure 5). BEN circuit dissection requires immobile gel-embedded fish, and proper motion parallax re-creation in a virtual setting depends upon an immersive visual environment that is tuned to the exact statistics of tail motion. Our group is currently addressing this issue using detailed Bayesian tail models that instantly and accurately update a 3D virtual world. This approach will create accurate motion parallax in an embedded setting, allowing the type of circuit dissection seen in previous studies to be conducted on zebrafish that are computing 3D object statistics.

## Materials and Methods

### Zebrafish

Experiments were conducted according to the guidelines of the National Institutes of Health and were approved by the Standing Committee on the Use of Animals in Research of Harvard University. All wild-type experiments were performed on dpf 7–9 larval zebrafish of the WIK strain (ZFIN ZDB-GENO-010531–2, https://zfin.org). For the Mauthner ablation experiments, a transgenic fish generated by our laboratory (MH 16946-Gal4, https://zebrafishexplorer.zib.de/home) was crossed to a UAS-Kaede reporter line (Tg(UAS-E1b:Kaede)s1999t; ZDB-ALT-070314-1, https://zfin.org), yielding green fluorescence in both Mauthner neurons. All fish were fed daily with paramecia starting at dpf4.

### Behavioral Setup

After a morning paramecia feeding, fish were added to a 12 cm-diameter acrylic tank with clear bottom and black walls. The tank was then mounted atop a clear acrylic stage. Fish behavior was recorded from above by a Mikrotron EoSens camera capturing at 500 Hz and 1280 × 1024 pixel resolution. Custom C# acquisition code was written for high-speed video writing, online background subtraction, stimulus delivery, and fish detection (available on Github). The entire rig was enclosed in black cardboard and tarp to minimize external light and noise, and all behavioral experiments were conducted in a dark room. A space heater was placed within the rig in order to maintain zebrafish at approximately 27° C. The tank was illuminated from below with an infrared light array to allow imaging in both light and dark conditions. Additionally, to draw fish into the center of the tank where barriers were located, a radial gradient phototaxis stimulus was projected onto a cold mirror and reflected onto the bottom of the tank. Once fish reached a threshold distance from the tank center, projector illumination was switched to bright whole-field light gray, which strongly lit the walls of the tank enclosure, reflecting light both onto the bottom and top of the tank. In dark trials, the projector was instead turned to whole-field black and automatically covered with a filter-flipper to block all light. Thus all escape stimuli were delivered under bright whole-field gray illumination or darkness (except one condition described in Figure 3). For experiments with physical barriers, circular pieces of extruded acrylic were affixed to the bottom of the dish with inert dental wax. Unless explicitly described in the text as white, barriers were dark red in color. In virtual barrier experiments, circles the same width and color as the smallest physical barriers (12 mm diameter) were projected onto the floor of the tank. Throughout the entire trial, the acquisition program logged the XY coordinates of the fish, as well as the coordinates and dimensions of the barriers. If the fish came within the trial’s threshold distance of a barrier edge, a stimulus command was sent via a PyBoard microcontroller to an electromagnet affixed to the stage. The electromagnet drives a metal rod into the stage for 200 ms, inducing a non-directional auditory startle stimulus (a “tap”) [22]. An LED light was fixed to the top of the metal rod such that its reflection into a mirror situated atop the stage was captured by the camera. From the LED reflection’s movement across the mirror, the exact timing of the tap was extracted at 2-millisecond resolution (b/c 500Hz acquisition) by Python-based analysis code (available on Github). There were two metrics for excluding fish in our assay: fish were excluded from analysis if they failed to enter the barrier zone for 5 minutes, 5 times, before completing 5 escape trials (20.2% of total tested fish). Second, we excluded fish if the LED indicating stimulus delivery indicated that the stimulus was not evoked. Lastly, because we excluded the frontal-most visual field (Figure 1, 3) from analysis between barrier conditions, we required that fish must have performed at least 3 trials in the given condition to be given PI or BAI score.

#### Snell’s Window

For all solid barrier conditions, the top of the barrier protruded above the water surface. By calculating apparent height due to refraction and checking for the collapse of the image into the horizon of Snell’s Window [12], we confirm that the tops of all tested barriers are visible to the fish, and that distortion of the barriers at the distances tested is minimal (the image formed by a 4 mm distant barrier of 6 mm height occupies 97.5% of the retinal height subtended by the 8 mm distant barriers at 12 mm high). Water refractive index used in our calculations was 1.333 and depth of the fish eye underwater 1 mm.

#### Escape Analysis

Upon receiving a tap stimulus, all fish engaged in fast escape swims characterized by initial high angle turns and a subsequent burst of speed (Supplementary Figure 1). To uncover the tail angle and direction of escape trajectories relative to barriers, 500Hz videos of escape sequences were background subtracted and dynamically thresholded using an OpenCV-based recursive contour-finding algorithm that terminated after finding a fish-sized object. The heading angle was recorded by calculating a vector arising from the midpoint of the fish contour and extending to the center of mass of the contour, which pinpoints the fish’s head. A vector from the fish contour to the nearest barrier was also calculated per trial; comparing this vector to the fish’s heading angle allowed us to detect whether the fish approached the barrier from the left or the right.

To analyze escape direction, we used the calculated heading angle averaged over the 20 ms before tap stimulus command to rotate the XY coordinates of the escape trajectory (20ms -¿ 60ms after tap command) so that escape coordinates are calculated with the fish initiated vertically and its origin at point (0, 0). In this way, positive X coordinates reflect escapes to the right and negative X coordinates reflect escapes to the left. We considered an escape to be “left” if the sum of escape X coordinates was negative and “right” if positive; we considered the escape an “avoidance” if the angle to the barrier relative to the heading angle of the fish was opposite in sign to the X coordinate sum. If the fish collided with the barrier during its escape trajectory (determined when the center of mass is within 750 microns of the barrier edge), the direction of escape was calculated only up until the collision timepoint and a collision was registered. This prevented the assignment of “avoidance” to trajectories where fish deflect off of barriers.

For tail angle calculation (Supplementary Figure 1), the tail was split into 6 segments and cumulative angles were calculated using dot products between segment vectors with direction determined by the cross product. The timeseries of tail angle sums was filtered with a Gaussian (=1). The first time after stimulus delivery that the relative maximum or minimum of the filtered timeseries passed + / - 30 degrees was demarcated as the escape initiation time, with its magnitude recorded (Supplementary Figure 1). Note that our characterization determines latency as the time from stimulus delivery until maximum tail amplitude is reached, not until the initiation of the escape as in other reports, meaning our latencies are slightly longer than typically reported.

#### Navigation Assay

To record the fish’s natural responses to obstacles while navigating, larvae that had never experienced taps in the vicinity of barriers were subject to the same experimental paradigm as above (e.g. phototaxis to attract the fish to the center, whole-field gray projection once barriers have been reached), except no taps were delivered when fish neared a barrier. The XY position of the fish and barriers, and the dimensions of the barriers, were tracked and plotted in Figure 1; for heatmaps reflecting the fish’s preferred swimming distance with respect to barriers, XY coordinates of the fish for every time point during navigation were calculated with respect to the nearest barrier. The coordinate system’s origin is therefore set as the nearest barrier center over time. Visited pixel locations were binned by Gaussian filtering (= 5) for the heatmap matrix representing the coordinate system.

### Mauthner Neuron Ablations

Phenylthiourea (PTU) treatment inhibits melanogenesis only over the duration of treatment so melanogenesis resumes on removal from the solution (Karlsson et al., 2001; Whittaker, 1966); we used PTU on fish before laser ablation to avoid damage caused by laser heating of melanocytes. Tg (MH 16946-Gal4 x UAS-Kaede) embryos were collected into 100 M phenylthiourea (PTU) embryo water solution. Larvae were allowed to develop in PTU solution starting at around 18 hpf until they were removed for ablation at 4 dpf. Larvae were returned to filtered fish water post-ablation and allowed to recover for two days. In practice, we found that PTU treatment decreased pigment expression but did not fully abolish pigmentation. Pigmentation increased upon removal from solution; this was necessary for detection of the tail during escape behavior. PTU treated larvae were screened on 4 dpf for minimal pigmentation and GFP expression in both Mauthner cells. Only GFP positive larvae in which both Mauthner cells were visible were used for ablation. Larvae were then embedded in 1.8 % low melting point agarose in the center of a petri dish. Mauthner neurons were localized under two-photon excitation. Laser pulsing at 800-850nm at 360 mW for .1-.3 seconds was used in 5-10 locations on the Mauthner neuron. This induced Mauthner cell explosion in a subset of fish; gross morphological deformation was visible under 950 nm illumination after successful ablations, confirming that disappearance of the cell was not due to photobleaching. Any larvae where clear explosion of the Mauthner did not occur after multiple ablation attempts were discarded. Larvae were then freed from agarose and allowed to recover for two days. They were fed with paramecia on the first day post-ablation (6 dpf) and behaviorally assayed from 7-9 dpf as described above.

## Data Availability

The data and code for analysis used in this study are openly available on Github which also includes the barrier locations, sizes, and raw escape trajectories for each trial performed by all fish.

The authors would like to thank Martin Haesemeyer, Josua Jordi, Rob Johnson, Eva Naumann, Tim Dunn, Kirsten Bolton, Kristian Herrera, Clemens Riegler, and Armin Bahl for discussions concerning experimental design and interpretation. Alix Lacoste’s excellent thesis defense was especially influential in laying the foundations of this project. All members of the Engert lab were instrumental in weekly discussions. This work was funded by a U19 grant from the NIH.

## Notes

### Competing Interest Statement

The authors have declared no competing interest.

## References

1. M. B. Ahrens, J. M. Li, M. B. Orger, D. N. Robson, A. F. Schier, F. Engert, and R. Portugues. Brain-wide neuronal dynamics during motor adaptation in zebrafish. Nature, 485(7399):471–477, May 2012.

2. S. Alem, C. J. Perry, X. Zhu, O. J. Loukola, T. Ingraham, E. Søvik, and L. Chittka. Associative Mechanisms Allow for Social Learning and Cultural Transmission of String Pulling in an Insect. PLOS Biology, 14(10):e1002564, Oct. 2016. Publisher: Public Library of Science.

3. A. J. Barker and H. Baier. Sensorimotor decision making in the zebrafish tectum. Current biology: CB, 25(21):2804–2814, Nov. 2015.

4. P. W. Battaglia, J. B. Hamrick, and J. B. Tenenbaum. Simulation as an engine of physical scene understanding. Proceedings of the National Academy of Sciences, 110(45):18327–18332, Nov. 2013. Publisher: National Academy of Sciences Section: Biological Sciences.

5. I. Bianco, A. Kampff, and F. Engert. Prey Capture Behavior Evoked by Simple Visual Stimuli in Larval Zebrafish. Frontiers in Systems Neuroscience, 5:101, 2011.

6. A. D. Bolton, M. Haesemeyer, J. Jordi, U. Schaechtle, F. A. Saad, V. K. Mansinghka, J. B. Tenenbaum, and F. Engert. Elements of a stochastic 3D prediction engine in larval zebrafish prey capture. eLife, 8:e51975, Nov. 2019. Publisher: eLife Sciences Publications, Ltd.

7. V. Braitenberg. Vehicles: Experiments in Synthetic Psychology. MIT Press, Feb. 1986. Google-Books-ID: 7KkUAT q sQC.

8. R. A. Brooks. Intelligence without representation. Artificial Intelligence, 47(1):139–159, Jan. 1991.

9. A. B. Chen, D. Deb, A. Bahl, and F. Engert. Algorithms underlying flexible phototaxis in larval zebrafish. Journal of Experimental Biology, 224(10):jeb238386, May 2021.

10. C. Chiandetti and G. Vallortigara. Intuitive physical reasoning about occluded objects by inexperienced chicks. Proceedings of the Royal Society B: Biological Sciences, 278(1718):2621–2627, Sept. 2011. Number: 1718 Publisher: Royal Society.

11. I. Couzin and J. Krause. Self-Organization and Collective Behavior in Vertebrates. Advances in The Study of Behavior - ADVAN STUDY BEHAV, 32:1–75, Dec. 2003.

12. T. W. Dunn and J. E. Fitzgerald. Correcting for physical distortions in visual stimuli improves reproducibility in zebrafish neuroscience. eLife, 9:e53684, Mar. 2020. Publisher: eLife Sciences Publications, Ltd.

13. R. C. Eaton and D. S. Emberley. How stimulus direction determines the trajectory of the Mauthner-initiated escape response in a teleost fish. Journal of Experimental Biology, 161(1):469–487, Nov. 1991. Number: 1.

14. R. C. Eaton, R. K. K. Lee, and M. B. Foreman. The Mauthner cell and other identified neurons of the brainstem escape network of fish. Progress in Neurobiology, 63(4):467–485, Mar. 2001.

15. B. d. Gelder, M. Tamietto, G. v. Boxtel, R. Goebel, A. Sahraie, J. v. d. Stock, B. M. C. Stienen, L. Weiskrantz, and A. Pegna. Intact navigation skills after bilateral loss of striate cortex. Current Biology, 18(24):R1128–R1129, Dec. 2008. Publisher: Elsevier.

16. D. Ingle. Focal attention in the frog: behavioral and physiological correlates. Science, 188(4192):1033–1035, 1975. Publisher: American Association for the Advancement of Science.

17. D. J. Ingle and K. v. Hoff. Visually Elicited Evasive Behavior in Frogs. BioScience, 40(4):284–291, 1990. Number: 4 Publisher: [American Institute of Biological Sciences, Oxford University Press].

18. G. Johansson. VISUAL MOTION PERCEPTION. Scientific American, 232(6):76–89, 1975. Publisher: Scientific American, a division of Nature America, Inc.

19. M. Koyama, F. Minale, J. Shum, N. Nishimura, C. B. Schaffer, and J. R. Fetcho. A circuit motif in the zebrafish hindbrain for a two alternative behavioral choice to turn left or right. eLife, 5:e16808, Aug. 2016. Publisher: eLife Sciences Publications, Ltd.

20. R. J. Krauzlis, A. R. Bogadhi, J. P. Herman, and A. Bollimunta. Selective attention without a neocortex. Cortex, 102:161–175, May 2018.

21. R. J. Krauzlis, L. P. Lovejoy, and A. Zénon. Superior Colliculus and Visual Spatial Attention. Annual Review of Neuroscience, 36(1):165–182, 2013. Number: 1 eprint: https://doi.org/10.1146/annurev-neuro-062012-170249.

22. A. Lacoste, D. Schoppik, D. Robson, M. Haesemeyer, R. Portugues, J. Li, O. Randlett, C. Wee, F. Engert, and A. Schier. A Convergent and Essential Interneuron Pathway for Mauthner-Cell-Mediated Escapes. Current Biology, 25(11):1526–1534, June 2015.

23. G. W. Lindsay. Attention in Psychology, Neuroscience, and Machine Learning. Frontiers in Computational Neuroscience, 14:29, 2020.

24. A. Lock and T. Collett. The Three-Dimensional World of a Toad. Proceedings of the Royal Society of London. Series B, Biological Sciences, 206(1165):481–487, 1980. Publisher: The Royal Society.

25. H. R. Maturana and F. J. Varela. The tree of knowledge: The biological roots of human understanding. The tree of knowledge: The biological roots of human understanding. New Science Library/Shambhala Publications, Boston, MA, US, 1987. Pages: 263.

26. R. McIntosh, K. McClements, I. Schindler, T. Cassidy, D. Birchall, and A. Milner. Avoidance of obstacles in the absence of visual awareness. Proceedings of the Royal Society of London. Series B: Biological Sciences, 271(1534):15–20, Jan. 2004. Number: 1534 Publisher: Royal Society.

27. M. Minsky. Society Of Mind. Simon and Schuster, Mar. 1988. Google-Books-ID: bLDLllfRpdkC.

28. M. Minsky. The emotion machine: Commonsense thinking, artificial intelligence, and the future of the human mind. Simon and Schuster, 2007.

29. D. M. O’Malley, Y.-H. Kao, and J. R. Fetcho. Imaging the Functional Organization of Zebrafish Hindbrain Segments during Escape Behaviors. Neuron, 17(6):1145–1155, Dec. 1996. Number: 6.

30. B. W. Patterson, A. O. Abraham, M. A. MacIver, and D. L. McLean. Visually guided gradation of prey capture movements in larval zebrafish. Journal of Experimental Biology, 216(16):3071–3083, 2013. Publisher: Company of Biologists.

31. M. Poteser and K. Kral. Visual Distance Discrimination Between Stationary Targets in Praying Mantis: An Index of the use of Motion Parallax. Journal of Experimental Biology, pages 2127–2137, 1995.

32. S. Schuster, R. Strauss, and K. G. Götz. Virtual-Reality Techniques Resolve the Visual Cues Used by Fruit Flies to Evaluate Object Distances. Current Biology, 12(18):1591–1594, Sept. 2002.

33. E. C. Sobel. The locust’s use of motion parallax to measure distance. Journal of Comparative Physiology A, 167(5):579–588, Nov. 1990.

34. E. S. Spelke and S. A. Lee. Core systems of geometry in animal minds. Philosophical Transactions of the Royal Society B: Biological Sciences, 367(1603):2784–2793, Oct. 2012. Number: 1603 Publisher: Royal Society.

35. M. V. Srinivasan. Distance Perception in Insects. Current Directions in Psychological Science, 1(1):22–26, Feb. 1992. Publisher: SAGE Publications Inc.

36. T. D. Ullman, E. Spelke, P. Battaglia, and J. B. Tenenbaum. Mind Games: Game Engines as an Architecture for Intuitive Physics. Trends in Cognitive Sciences, 21(9):649–665, Sept. 2017. Publisher: Elsevier.

37. G. K. Wallace. Visual Scanning in the Desert Locust Schistocerca Gregaria Forskål. Journal of Experimental Biology, 36(3):512–525, Sept. 1959. Number: 3.

