## Supplemental Figure 1 for "Covert attention to obstacles biases zebrafish escape direction"

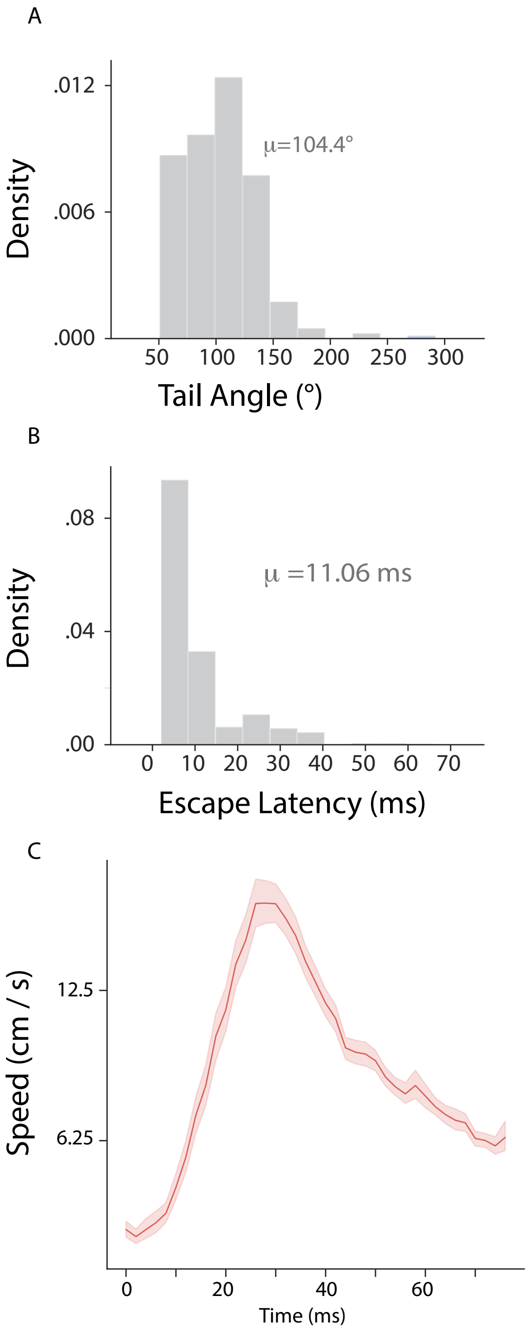


Figure S**1:** A**.** Distribution of maximum cumulative tail angle per escape for all fish tested in barrier conditions**.** B**.** Distribution of escape latencies **(**ms from tap contact with dish until maximum tail angle**).** C**.** Timeseries plot of velocity of the fish starting at tap command (light red = 95% CI)**.**
